# Senkyunolide A protects corticosterone-induced cell apoptosis via modulating protein phosphatase 2A and α-synuclein

**DOI:** 10.1101/145326

**Authors:** Shenglan Gong, Jin Zhang, Zhouke Guo, Wenjun Fu

## Abstract

*Dan-zhi-xiao-yao-san* is a Traditional Chinese Medicine (TCM) formula that is widely used to treat depression related neurological disorders, however, the active compound(s) and underlying mechanisms are unclear. In the present study, we found that senkyunolide A (SenA) has neuroprotective effects in corticosterone (Cort)-induced depression cell model in PC12 cells. Firstly, we found that SenA protects Cort-induced cell injury in PC12 cells. In addition, SenA attenuates Cort-induced the reduction of phosphatase 2A (PP2A) activities, and the increase of p-PP2A, α-syn and p-α-syn-Ser129 levels. Furthermore, PP2A inhibitor okadaic acid (OA) decreased, and PP2A activator D-erythro-Sphingosine (SPH) increased Cort-induced cell apoptosis. Importantly, we also found that the neuroprotective effects of SenA in Cort-induced cell injury via modulating α-syn levels. Collectively, our results suggest that the neuroprotective effects of SenA in Cort-induced depression cell model via modulating PP2A activities and α-syn levels, and bring a breakthrough to the anti-depression mechanisms for natural compound SenA.

## Introduction

Depression is one of the most common life-threatening psychiatric disorders and has a high prevalence[1, 2]. Though the pathogenesis of depression is not well characterized, hyperactivation of the hypothalamic-pituitary-adrenal (HPA) axis has been implicated in the pathogenesis and progression of this disease[2–4]. Hyperactivation of HPA is characterized by higher levels glucocorticoids in the circulating blood[5, 6]. Elevated concentrations of blood glucocorticoid are found in depression patients[2, 4, 5]. In addition, high glucocorticoid levels cause depression-like behavior in animals and induce pathological injury to the hippocampal neurons both *in vitro* and *in vivo*[2, 6]. The rat pheochromocytoma (PC12) cell line, express relative high levels of glucocorticoid receptors and has typical neuron features, is one of widely used neuronal cell lines for neuroscience related studies[7]. High concentrations of glucocorticoid induced PC12 cell injury has been using as an effective in vitro experimental model for depression study[8, 9]. Importantly, a variety of classic antidepressants have been shown to have cytoprotective effects in this cell model[10,11], suggesting that antidepressants may act by protection against glucocorticoid-induced neurotoxicity.

Currently, though several Western drugs, such as monoamine oxidase inhibitor, tricyclic antidepressants, selective serotonin reuptake inhibitor, noradrenaline reuptake inhibitor and noradrenergic reuptake inhibitor are commonly clinical application[12–14]. However, seriously side effects hurdle their applications[15, 16]. Consequently, developing for a better-tolerated, safer and powerful antidepressant is an urgent need. Traditional Chinese Medicine (TCM) has been used for thousands of years for treating of depression related diseases in China and other Asia countries. Thus, discovering of novel anti-depression agents from TCM may be a promising alterative strategy. *Xiao-yao-san* and modified formula such as *Dan-zhi-xiao-yao-san* are most commonly used TCM formulas for treating multiple diseases such as depression in China and other Asia countries[17]. Based on TCM theory, *Xiao-yan-san* mainly used for promoting liver qi circulation, coursing the liver, nourishing liver blood and fortify the spleen[18–20]. *Dan-zhi-xiao-yao-san* is comprised of several herbs including *atractylodismacrocephalerhizoma*, *bupleuri radix*, *angelicaesinensis*, *poria*, *glycyrrihizae radix*, *tree peony bark*, *gardenia jasminoides*, *paeonialactiflora pall*, *mint* and *roasted ginger*[18–20]. Although *xiao-yao-san* is one of the mostly commonly used formulas in China for treating depression, the active compound(s) and underlying mechanism are largely unknown.

In the present study, we firstly tested the neuroprotective effect of several major compounds in *Dan-zhi-xiao-yao-san* in corticosterone-induced depression cell model and found that senkyunolide A is the most effective one. We next investigated the underlying mechanisms focusing on synuclein and PP2A pathways.

## Materials and methods

### Cell culture

PC12 cells were obtained from the American Type Cell Culture Collection (Manassas, VA, USA) and were maintained in Dulbecco’s modified Eagle’s medium (DMEM) supplemented with 10% horse serum and 5% fetal bovine serum (FBS), and 1% penicillin/streptomycin at 37 °Cin a humidified 5% CO_2_ atmosphere.

### Cell viability assay

The cell viability was evaluated by CCK-8 assay (Dojindo Molecular Technologies,Inc.). Briefly, PC12 cells were plated in the 96-well plates, after 24 h, cells were treated with different drug for another 24 h, then washed with D-Hanks buffer solution. Finally, 200 µlCCK-8 solution was added to each well and incubated for an additional 3 h at 37 ◦C. The optical density (OD) of each well at 450nm was recorded on a Microplate Reader (Thermo, Varioskan Flash).

### Lactate dehydrogenase (LDH) leakage assay

The release of LDH is amarker for cellular toxicity. LDH activity was measured using a LDH diagnostic kit (from Thermo Fisher, 88953) according to the manufacturer’s instructions. Briefly, PC12 cells were seeded in 24-well culture plates at a density of 1×105 cells/well. After treatment with SenA in the presence or absence of Cort, the medium was collected for analysis of LDH activities.

### Flow cytometric analysis of cell apoptosis

At the end of drug treatment, cells were freshly harvested and suspended in a 1 : 1 (v/v) mixture of PBS and 0.2 M Na_2_HPO_4_-0.1 M citric acid (pH 7.5). Following the fixation with ice-cold ethanol at 4 °C for 1 h, the cells were resuspended in binding buffer and then incubated in a buffer containing 200ng/ml Annexin V-FITC conjugates (Sigma, MO, USA) at room temperature for 15 min. Subsequently, the cells were stained with PI (Sigma, MO, USA) (300 ng/ml) for 10 min. The stained cells were analyzed on a FACSCalibur flow cytometer (BD Biosciences, USA).

### Immunofluorescence staining

At the end of drug treatment, cells were fixed with 4% paraformaldehyde in PBS, permeabilized with 0.5% Triton X-100 and then blocked with 5% normal goat serum. The cells were then incubated with anti-α-syn (Abcam, UK) and anti-PP2A (Abcam, UK), antibodies in 1% BSA at 4 ˚C overnight. Cell nuclei were indicated by staining with 4’-6-diamidino-2-phenylindole (DAPI) for 15 minutes. The immunofluorescence images were acquired on Zeiss fluorescence microscopy (Carl Zeiss, Germany).

### Western blotting analysis

Cells were harvest by RIPA Lysis and Extraction Buffer (89900, Thermo Fisher Scientific). Denatured protein samples were resolved on SDS-PAGE and transferred to PVDF membrane (Millipore, Billerica, MA, USA). After blocking with non-fat milk, membrane was incubated overnight at 4 C with antibodies including α-syn (Abcam, UK), p-α-syn (Abcam, UK), PP2A (Abcam, UK) and GAPDH (Abcam, UK), followed by incubation with the anti-rabbit HRP-conjugated secondary antibodies (Abcam, UK). Chemiluminescence detection was performed using ECL advance Western blotting detection reagents (GE healthcare, Little Chalfont, Buckinghamshire, UK).

### Real-time PCR analysis

PC12 cells were used to analyze the expression of α-syn, PP2A and internal control β-actin by quantitative real-time PCR. Briefly, total RNA was extracted using TRIzol reagent (Invitrogen, CA, USA), and used for cDNA synthesis with 2 µg RNA and a High-Capacity cDNA Reverse Transcription kit. The expression of these three genes was conducted by real-time PCR using a kit from (DBI, German).The expression level of genes was calculated with normalization to housekeeping gene β-actin. Three independent experiments were carried out and each experiment performed in three analytical replicates. The forward and reverse primers for α-syn are showed in Table 1.

**Table 1.**
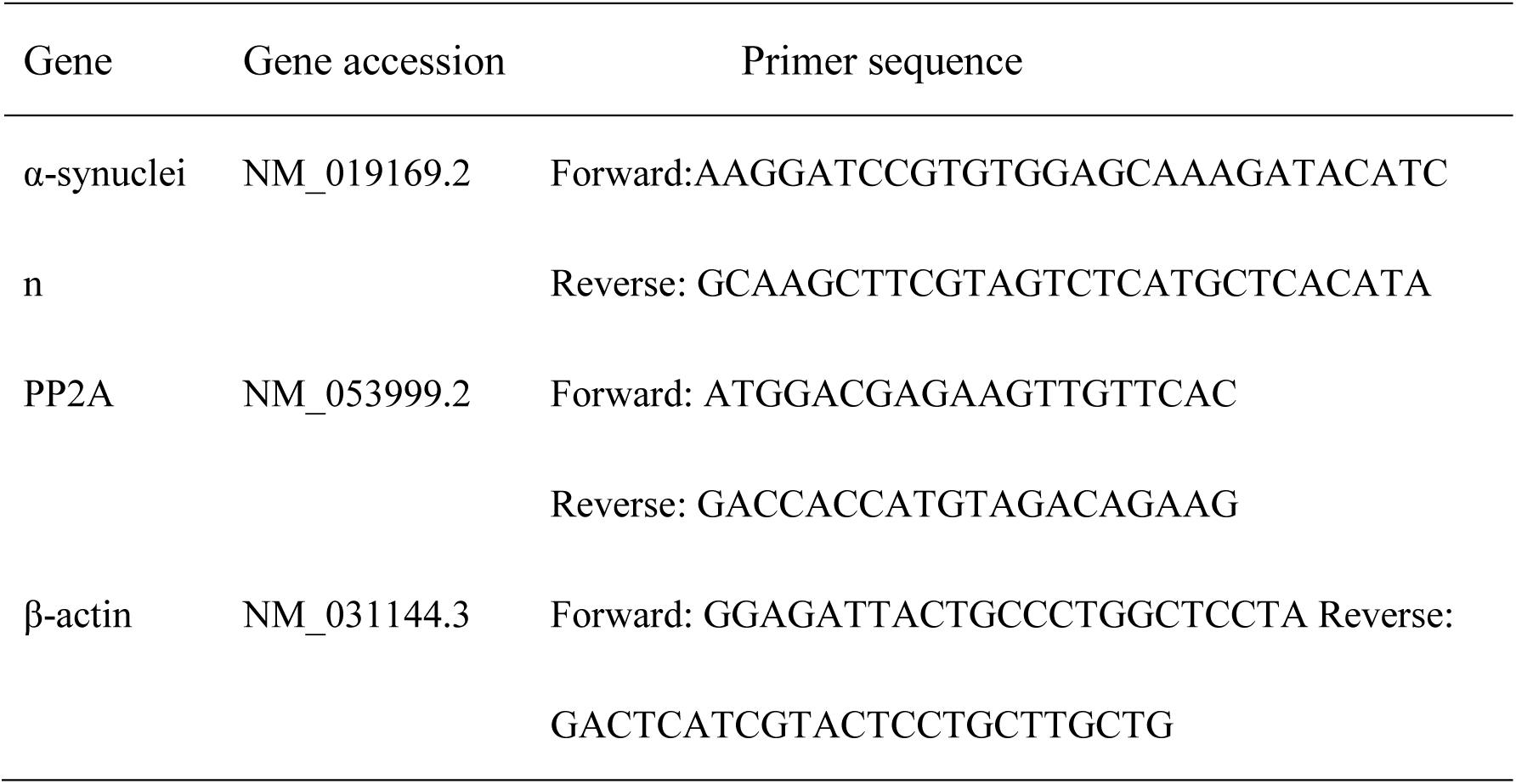
α-synuclein, PP2A and β-actin primers.

### Determination of PP2A activities by ELISA assay

PP2A activities was detectedby ELISA was used as described previously [21]. Briefly, celllysis buffer with Triton X-100 and protease inhibitors (10 μg/ml Aprotinin, 10 μg/mL Leupeptin, 100 μM PMSF, and 10 μg/ml Pepstatin A) were used for cell homogenization. Homogenized cell samples were centrifuged at 10,000 X g for 10 min at 4°C. Supernatant was collected for the protein phosphatase assay. The *pNPP* (*p*-Nitrophenyl phosphate) is a colorimetric substrate used for measuring the activity of serine/threonine phosphatases (Maehama et al. 2000). Assay buffer for PP2A (40 mM Tris-HCl, pH 8.4, 34mM MgCl2, 4mM EDTA and 4mM DTT). Upon dephosphorylation by phosphatases, *pNPP* turns yellow and read at absorbance 405 nm.

### Cell transfection

Αlpha-syn knockdown experiment was transfeted with siRNA with Lipofectamine^®^ RNAiMAX Reagent, over-expression experiment was transfected with Lipofectamine^®^ P3000 Reagent from Thermo Fisher Scientific. After 72 h of transfection, cells were used for experiments.

### Statistics analysis

Data were expressed as means ± standard deprivation (SD). Statistically significant differences between the groups were identified by one-way ANOVA analysis followed by paired Student’s t-tests. Differences at *P<0.05* and *P<0.01* were considered statistically significant.

## Results

### Neuroprotective effects of SenA against Cort-induced cytotoxicity

*Dan-zhi-xiao-yao-san* formula is one of most commonly used prescription for treating depression in China[22]. However, the activity compound(s) and underlying mechanisms are unclear. To determine which compound may contribute the anti-depression effects, we tested the neuroprotective effects of several major compounds in *Dan-zhi-xiao-yao-san* formula by using Cort-induced cell injury in PC12 cells as an in vitro cell model. By treating cells with geniposide, paeonol, ferulic acid, paeoniflorin, and SenA at the concentrations ranging from 0 to 2mg/ml for 24, 48, and 72 h, as shown in Figure 1A–E, SenA at the concentrations of 0.125 to 0.5 mg/ml significantly reversed Cort-induced cell viability in a time- and dose-dependent manner as reflected by CCK-8 assay (Figure 1E). However, for SenA at 2 mg/ml, a clearly cytotoxicity on PC12 cells was observed, which indicated that application of SenA in practice should be carefully managed. The chemical structure of SenA was shown in Figure 2A. Importantly, we found that SenA at 0.5 mg/ml concentration is the most effective concentrations possess neuroprotective effects, consequently, we chose this concentration for following mechanism studies.

**Figure 1.**
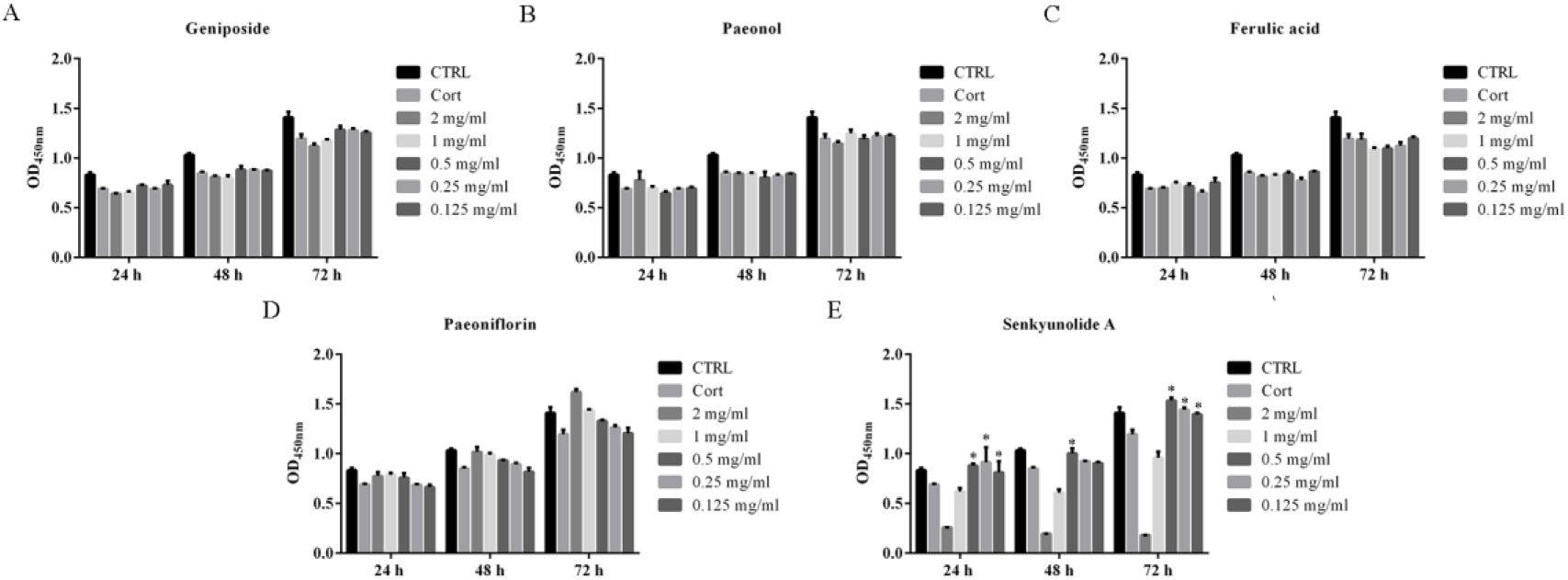
Neuroprotective effects of main compounds in Dan-zhi-xiao-yan-san in Cort-induced cell apoptosis in PC12 cells. (**A-E**) CCK-8assay determined the neuroprotective effects of geniposide, paeonol, ferulic acid, paeoniflorin, and SenA for Cort-induced cell death at the concentrations ranging from 0 to 2 mg/ml for 24, 48, and 72 h.*p<0.05 *vs* Cort group.

**Figure 2.**
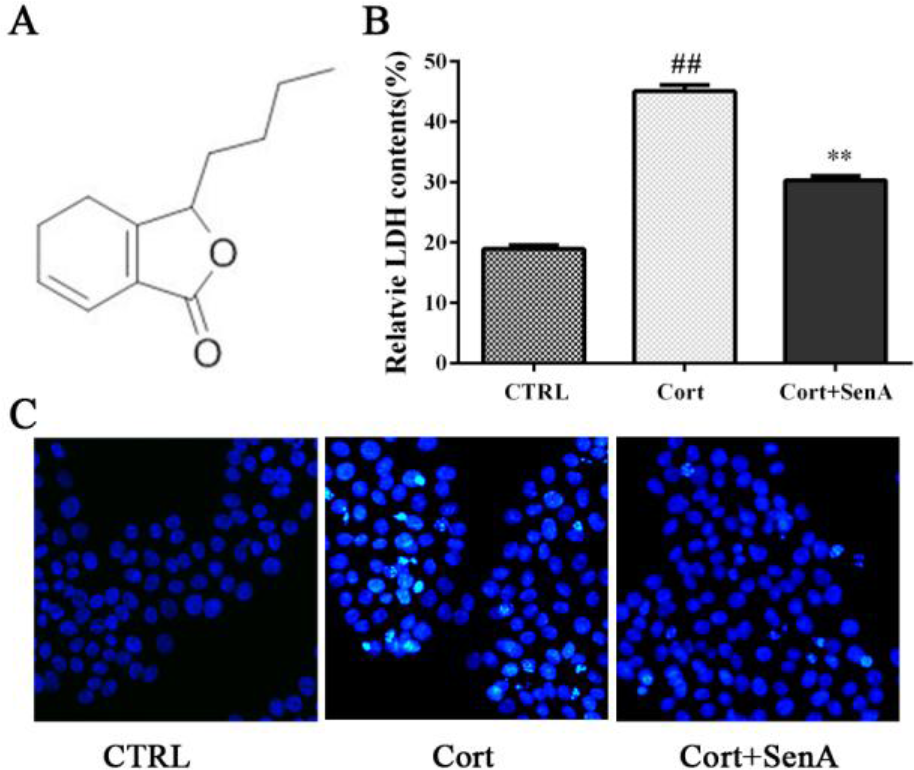
Neuroprotective effects of Sen A in Cort-induced cell apoptosis in PC12 cells. (**A**) Chemical structure of SenA. (**B**) LDH.assay showed that SenA significantly attenuated Cort-induced cell death. (**C**) Hoechst staining results show that SenA significantly attenuated Cort-induced cell apoptosis. ##p<0.01*vs* CTRL group, **p<0.01*vs* Cort group.

To further confirm the neuroprotective effects of SenA against Cort-induced cell injury. We next measured the effects of SenA against Cort-induced LDH release. LDH is a soluble cytosolic enzyme, which is widely presented in eukaryotic cells[23]. After cell death or necrosis, LDH is released to cell culture medium due to the damage of plasma membrane. Thus, the levels of LDH in cell culture medium is normally used as a marker for cell injury[23]. As shown in Figure 2B, the released LDH was significantly increased in Cort treated cells as compared with the control group. In contrast, pretreatment the cells with 0.5 mg/ml of SenA effectively decreased Cort-induced LDH release. In addition, Hoechst staining also showed that SenA can protect Cort-induced cell apoptosis (Figure 2C), Finally, Annexin V-FITC/PI staining results further support the notion that SenA has neuroprotective effects against Cort-induced cell apoptosis (Figure 3A–B).

**Figure 3.**
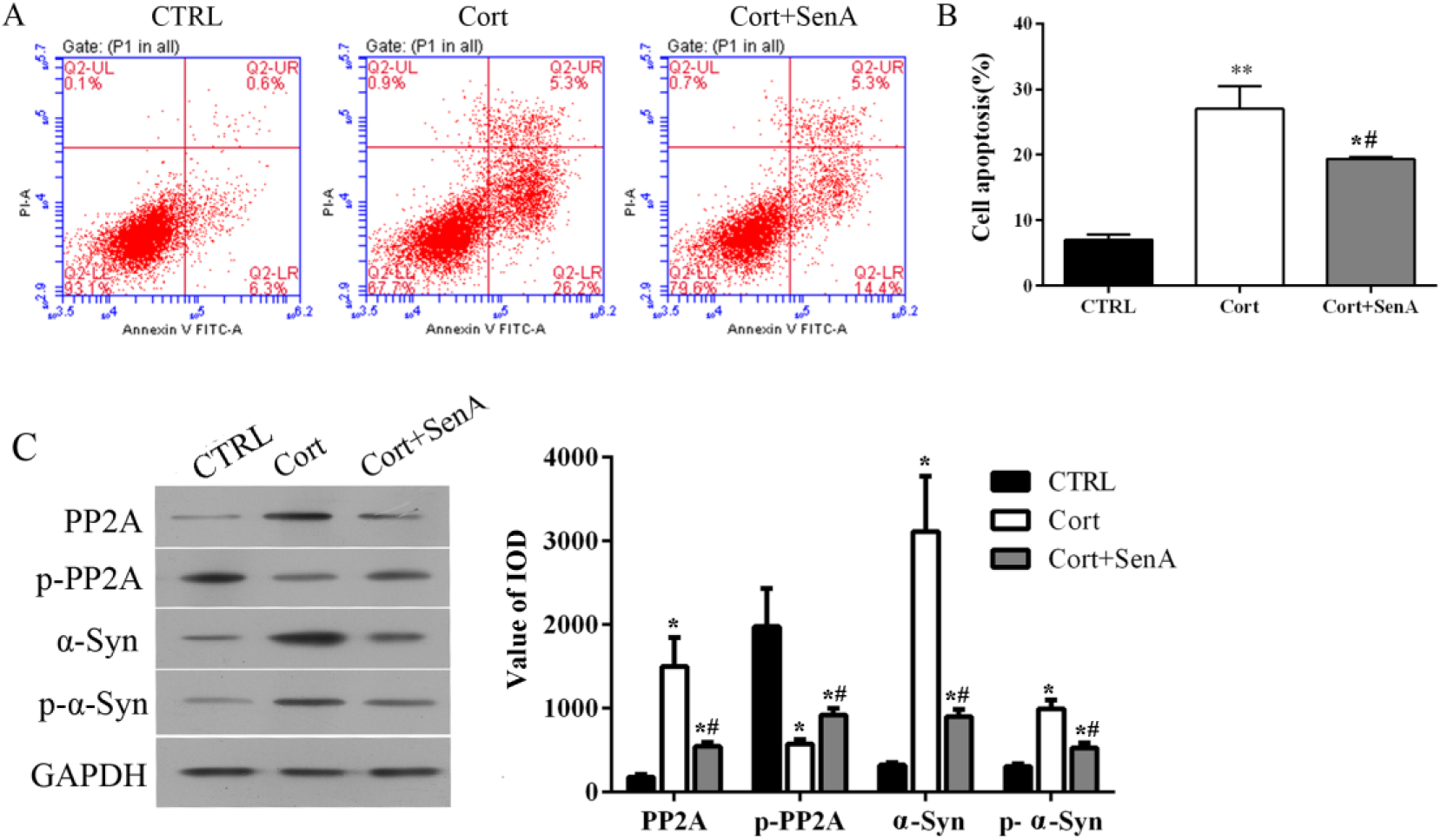
Involvement of PP2A and α-syn levels for the cytoprotective effect of SenA. (**A**) Annexin V-FITC/PI staining analysis showed that SenA significantly attenuated Cort-induced apoptosis. (**B**) Quantification data in A. (**C**) Western blotting results show that SenA attenuates Cort-induced the changes in the expression of PP2A, p-PP2A, α-syn and p-α-syn levels. *p<0.05 *vs* CTRL group, **p<0.01*vs* CTRL group. #p<0.0 *vs* Cort group.

### Involvement of the changes of PP2A and α-syn levels for the cytoprotective effect of SenA

Recent study revealed that over-expression of α-syn in neurons results in depression like symptoms in an animal model[24]. In addition, Cort up-regulates the expression of α-syn and the phospho-Ser129 α-syn is the most frequently modifier of α-syn[25]. Moreover, protein phosphatase 2A can regulate phospho-Ser129 α-synstates[26]. These results suggest that PP2A and α-syn are critical for depression. However, the neuroprotective effects of SenA via regulating these pathways are unclear. To examine whether PP2A and α-syn are involved for the cytoprotective effects of SenAin Cort-induced depression cell model. We found that Cort significantly increased the expression of α-syn and phosphorylation levels (Figure 3C). In contrast, SenA attenuates these effects (Figure 3C). In addition, α-syn is reported to reduce PP2A expression and activities. Our results demonstrated that SenA lowered Cort-induced the reduction of PP2A protein expression and phosphorylation levels (Figure 3C). PP2A activity assay by using ELISA further confirm this phenomenon (Figure 4E). Taken together, these results suggest that involvement of PP2A and α-syn for the cytoprotective effect of SenA in Cort-induced depression model.

**Figure 4.**
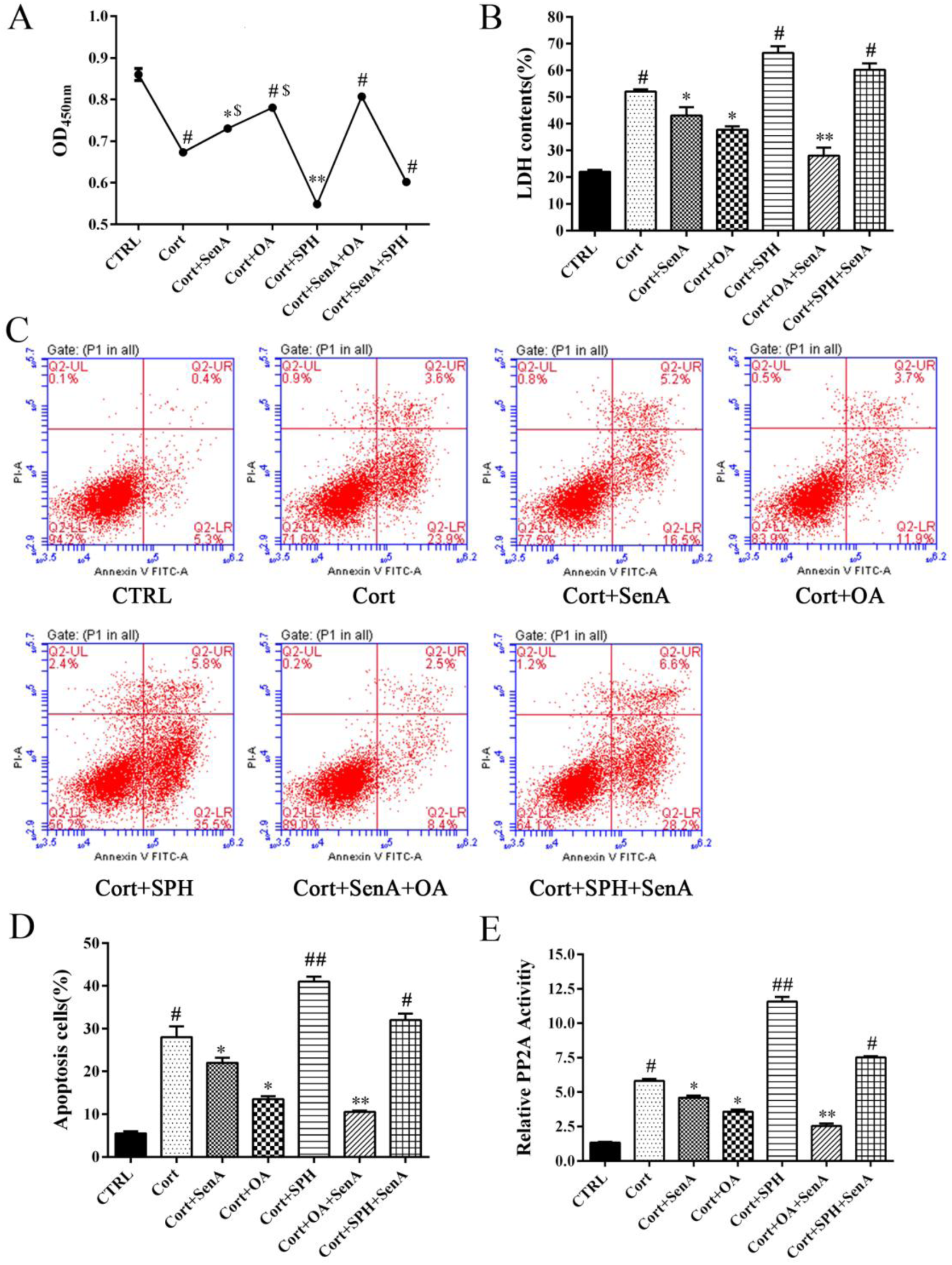
PP2A contribute the neuroprotective effects of SenA against Cort-induced cell injury. (**A**) CCK-8 assay show that PP2A inhibitor okadaic acid (OA) decreased, and PP2A activator D-erythro-Sphingosine (SPH) increased Cort-induced cell viability. SenA and PP2A activator SPH synergistically protect Cort-induced cell apoptosis. (**B**) LDH assay show that PP2A inhibitor okadaic acid (OA) increased, and PP2A activator D-erythro-Sphingosine (SPH) decreased Cort-induced cell injury. SenA and PP2A activator SPH synergistically protect Cort-induced cell apoptosis (p<0.05). (**C&D**) Annexin V-FITC/PI staining and quantification data further confirm the results in A&B. (**E**) ELISA assay determined PP2A protein activities after treated with PP2A inhibitor okadaic acid (OA) and PP2A activator D-erythro-Sphingosine (SPH). *p<0.05 *vs* Cort group. **p<0.01*vs* Cort group. #p<0.05 *vs* CTRL group. ##p<0.01*vs* CTRL group. $p<0.05 *vs* Cort+SenA+OA group.

### PP2A contribute the neuroprotective effects of SenA against Cort- induced cell injury

To further examine the roles of PP2A on regulation of SenA-mediated neuroprotective effects in Cort-induced depression cell model, we treated cells with PP2A inhibitor okadaic acid (OA) and PP2A activator D-erythro-Sphingosine (SPH) alone or in combination with SenA. As shown in Figure 4A, CCK-8 assay demonstrated that OA increased, while SPH decreased Cort-induced cell viability. SenA have synergistic effect with OA for neuroprotective effects in reducing LDH released caused by Cort (Figure 4B). Moreover, Annexin V-FITC/PI staining further confirmed that SenA have synergistic effect with OA for attenuating Cort-induced cell apoptosis (Figure 4C–D). As expected, PP2A inhibitor OA significantly inhibited PP2A activities as reflected by ELISA assay, whereas PP2A activator SPH enhanced PP2A activities (Figure 4E). Moreover, the results of immunofluorescence staining further confirmed the effects of OA decreased, and SPH increased the expression levels of PP2A levels (Figure 5). Taken together, these results suggest that activation of PP2A activities increases Cort-induced cell injury, whereas inhibition of PP2A attenuates Cort-induced cell injury. Remarkably, SenA and PP2A inhibitor OA can synergistically protect Cort-induced cell injury.

**Figure 5.**
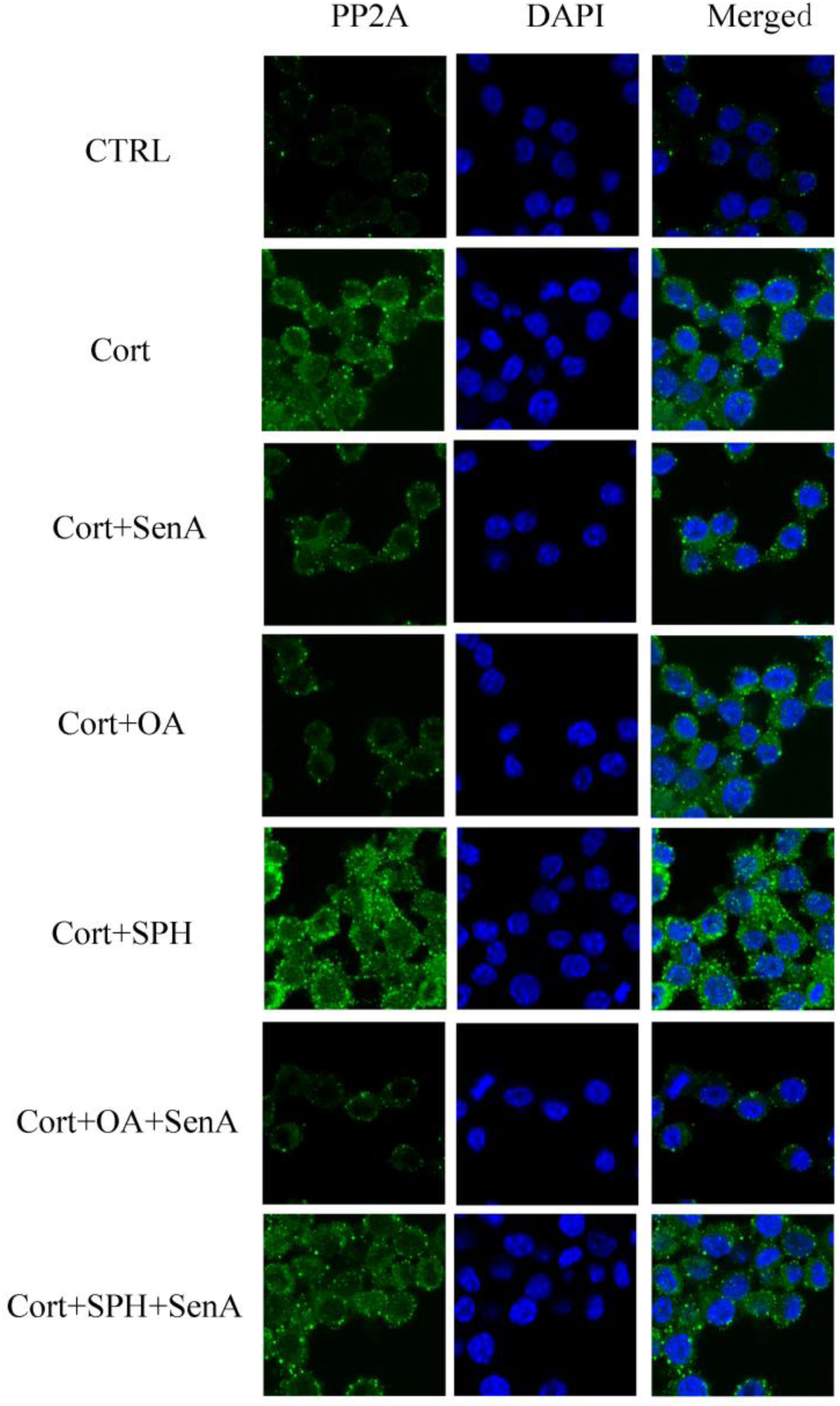
Effects of Sen A on the expression of PP2A levels after treated with PP2A inhibitor okadaic acid (OA) and PP2A activator D-erythro-Sphingosine (SPH). Immunostaining detected the expression of PP2A levels after treated with OA and SPH. SenA attenuated the Cort-induced the expression of PP2A levels, whereas OA reduces, whereas SPH increases PP2A levels after treated with Cort.

### α-syn contribute to neuroprotective effects of SenA against Cort-induced cell injury

As α-syn has been implicated in depression[24], we wonder whether α-syn and p-ser-129 phosho-α-syn contribute the neuroprotective effects of SenA in Cort-induced depression cell model. We firstly over-expressed and knocked down the expression of α-syn in PC12 cell, as shown in Figure 6A–C, transfection of cells with α-syn siRNA significantly decreased the expression of α-syn in both mRNA and protein levels. In addition, Western blotting and Q-PCR both showed that transfection of cells with α-syn plasmid significantly increased α-syn levels (Figure 6B–D). CCK-8 assay showed that knockdown the expression of α-syn increased cell viability, whereas over-expression of α-syn decreased cell viability (Figure 6E). Our LDH assay (Figure 6F), and Annexin V-FITC/PI assay (Figure 6G–H) further confirm these results. Importantly, Down-expression of α-syn significantly attenuates the neuroprotective effects of SenA in Cort-induced cellular damage (Figure 6E–H). Remarkably, α-syn overexpression does not further enhanced the neuroprotective effects of SenA in Cort-induced cell apoptosis, suggesting that SenA-induced neuroprotective effect may via down-regulation the expression of α-syn. Importantly, the immunofluorescence staining results further confirmed that knockdown the expression of α-syn decreased, whereas overexpression of α-syn increased Cort-induced the expression of α-syn levels (Figure 7).

**Figure 6.**
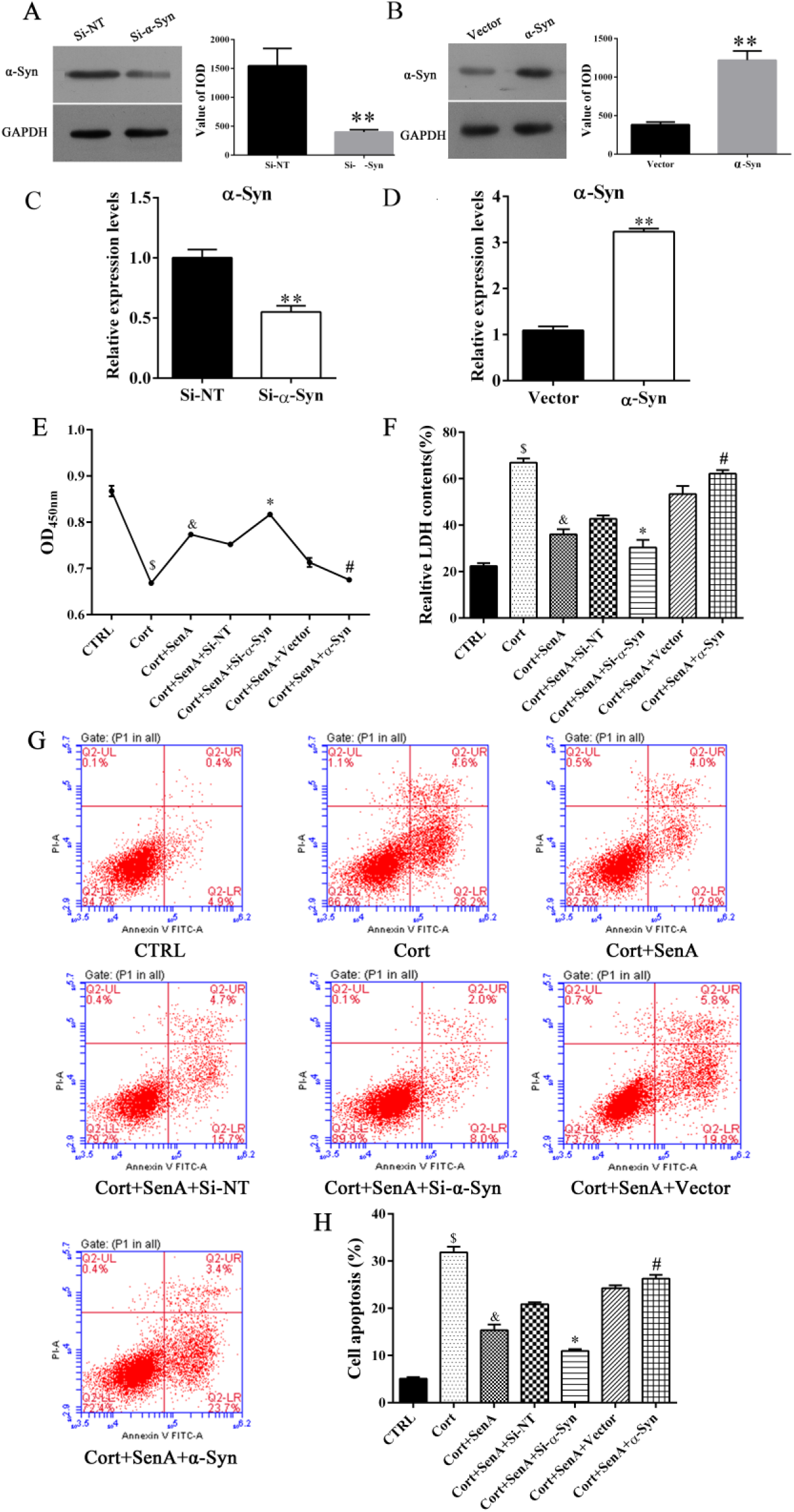
α-syn contribute the neuroprotective effects of SenA against Cort-induced cell injury. (**A**)Western blotting detected the expression of α-syn levels after transfected with α-syn specific siRNA. **p<0.01 (**B**) Western blotting detected the expression of α-syn levels after transfected with α-syn plasmid. **p<0.01. (**C&D**) Q-PCR indicates that transfection of α-synsiRNA decreased, whereas transfection of α-syn plasmid significantly increases the expression of α-syn levels.**p<0.01 (**E**) CCK-8assay revealed that α-syn overexpression decrease, α-syn knockdown decrease cell viability compare to CTRL group.*p<0.01 In contrast, Sen A cannot rescue cort-induced cell death in α-syn overexpressed cells. (**F**) LDH assay show similar results to E. (**G&H**) Annexin V-FITC/PI satining and quantification assay further confirmed that α-syn overexpression decrease, α-syn knockdown decrease cell apoptosis. In contrast, Sen A cannot rescue cort-induced cell apoptosis in α-syn overexpressed cells.*p<0.05 Cort+SenA+Si-α-Syn *vs* Cort+SenA group. #p<0.05 Cort+SenA+α-Syn *vs* Cort+SenA group. $p<0.05 *vs* CTRL. &p<0.05 *vs* Cort.

**Figure 7.**
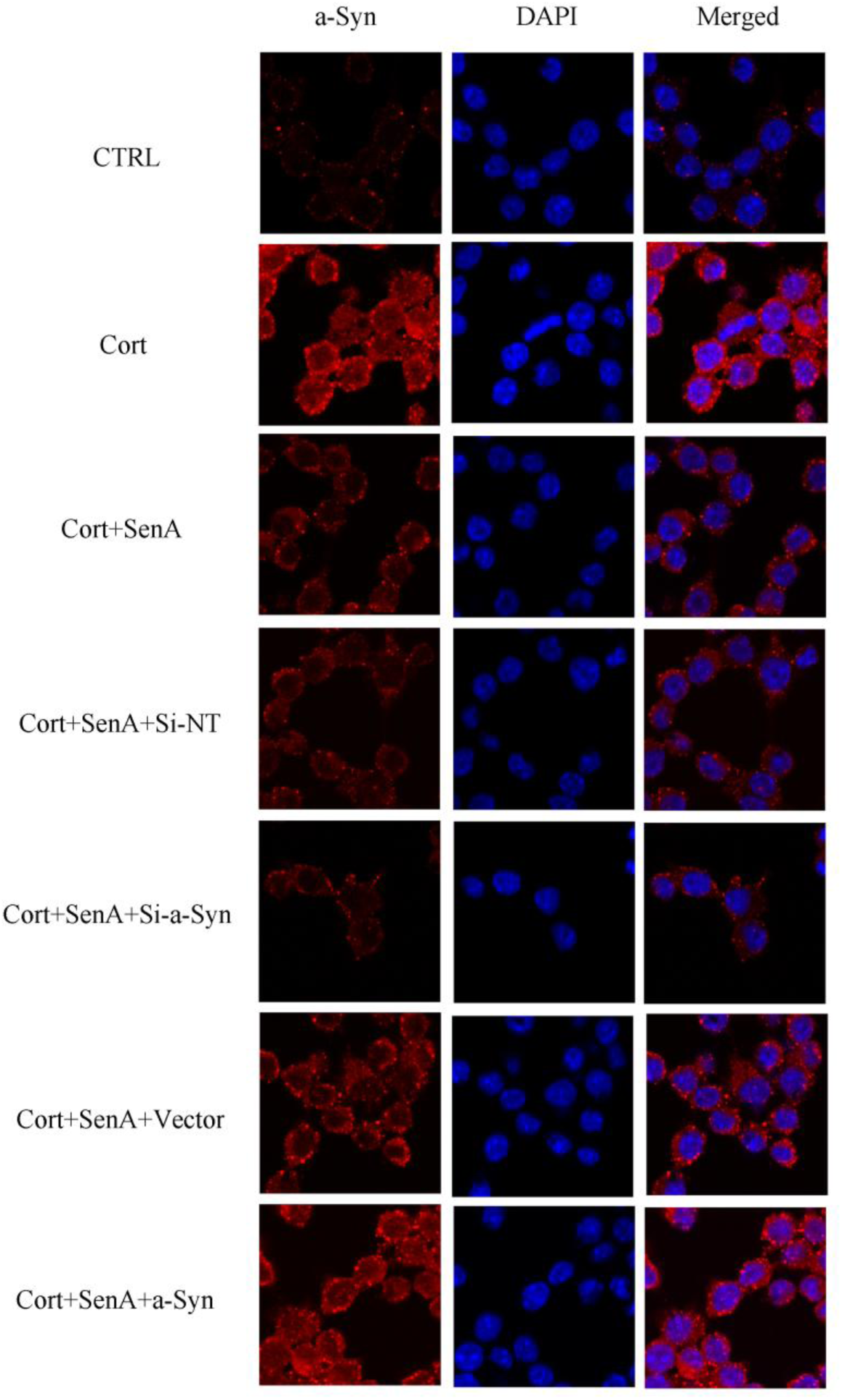
Immunostaining determined the expression of α-syn levels after knockdown or overexpression of α-syn. Immunostaining results shows that knockdown the expression of α-syn decreased, whereas overexpression of α-syn increased Cort-induced the expression of α-syn levels.

Finally, we examined the effects of the phosphorylation of α-syn (p-ser-129) on the neuroprotective effects of SenA against Cort-induced cell injury. We mutated Ser-129 to Ala-129 (SNCA-129A) to prevent the phosphorylation of Ser-129of α-syn. After transfection SNCA-129A plasmid into cells, western blotting results showed that the phosphorylation of α-syn significantly decreased phosphorylation states of α-syn at Ser-129 (Figure 8A). In addition, the mutation of α-syn Ser-129 to Ala-129 significantly attenuated Cort-induced the changes of cell viability as reflected by CCK-8 assay (Figure 8B). Moreover, LDH assay and Annexin V/FITC-PI assay further confirmed the protection effects of phosphorylation states of α-syn in Cort-induced cell injury (Figure 8C–E). Taken together, these results suggest that α-syn Ser-129 mutation to Ala-129 protect Cort-induced cell death. Moreover, the mutation of α-syn Ser-129 to Ala-129 also significantly decreased PP2A activities (Figure 8F). Taken together, these results suggest that the expression of α-syn levels and Ser-129 phosphorylation of α-syn both contribute to the neuroprotective effects of SenA in Cort-induced cell apoptosis.

**Figure 8.**
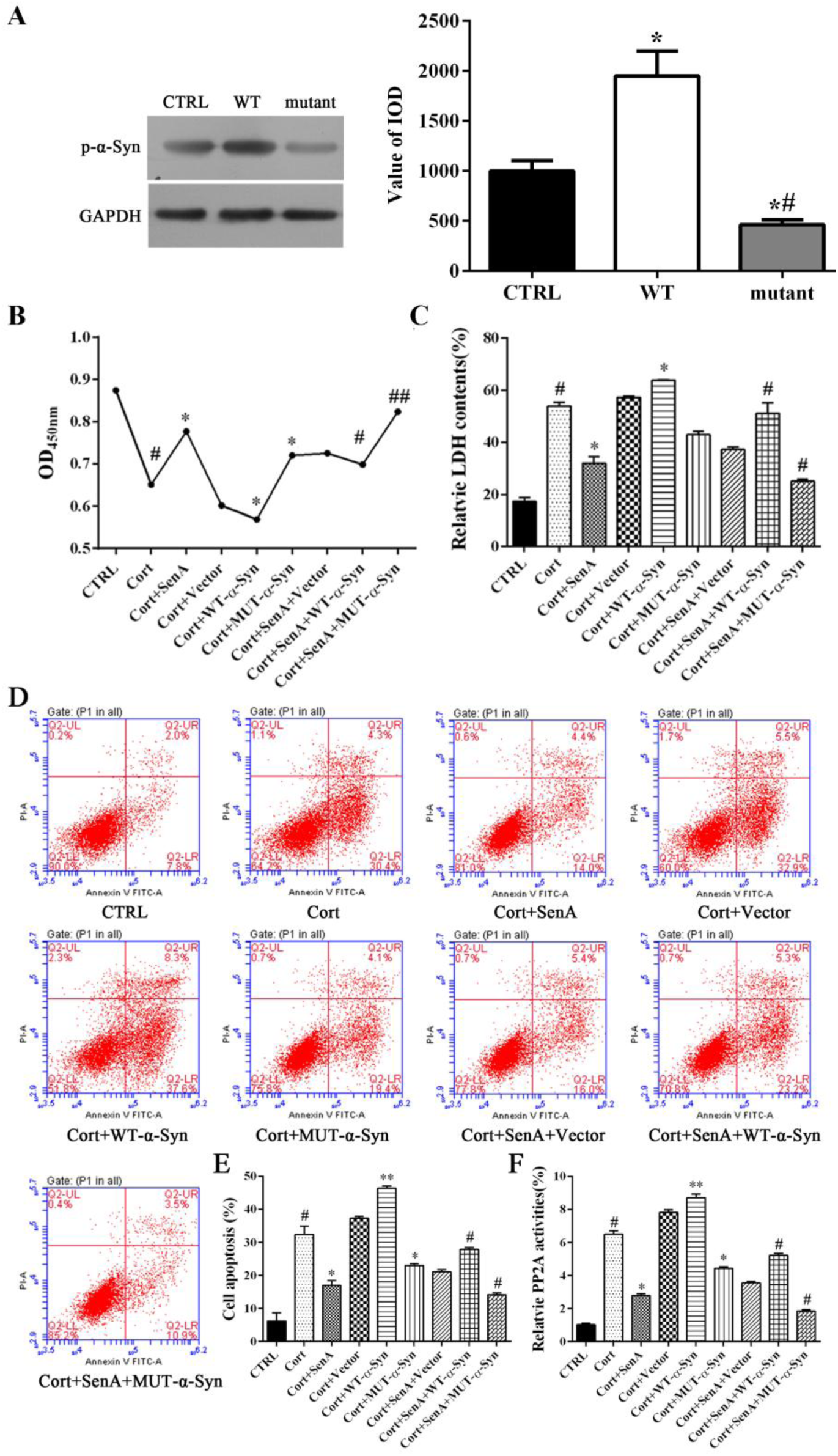
α-syn Ser129 phosphorylation is essential for the neuroprotective effects of SenA against Cort-induced cell injury. (**A**) After transfected cells with wild type (WT) and mutant (Ser 129 to Ala-129) α-syn, the expression of p-ser129 α-syn were detected. *p<0.05 *vs* CTRL group. **p<0.01 *vs* CTRL group. #p<0.0 *vs* Cort group. (**B**) CCK-8assay shows that WT α-syn increase Cort-induced cell injury, whereas Ala129 mutant and Sen A reduced Cort-induced cell injury. (**C**) LDHassay show similar results to B. (**D&E**) Annexin V-FITC/PI staining and quantification data further support that WT α-syn increase Cort-induced cell injury, whereas Ala-129 mutant and Sen A reduced Cort-induced cell apoptosis. (**F**) ELISA assay that WT α-syn increase Cort-induced PP2A activities, whereas Ala129 mutant and Sen A reduced Cort-induced PP2A activities. *p<0.05 *vs* Cort group. **p<0.01 *vs* Cort group. #p<0.05 *vs* Cort+SenA group. ##p<0.01 *vs* Cort+SenA group.

## Discussion

In the present study, we demonstrate that SenA shows neuroprotective effects in Cort-induced cell apoptosis in PC12 cells. The underlying mechanism may be through modulating PP2A activities and α-syn levels as well as α-syn’s phosphorylation.

*Xiao-yao-san* and its modified formulas such as *Dan-zhi-xiao-yao-san* are applied to treat a variety of diseases including menopausal syndrome, anemia, functional uterine bleeding, hepatitis, chronic gastritis, pelvic inflammatory disease and emotional diseases such as anxiety and depression [17, 22]. However, the active compound and underlying mechanisms for anti-depression effects are largely unclear. In the current study, we found that SenA is the active compound responsible for protection against Cort-induced cell apoptosis in PC12 cells among several tested compounds in such as ferulic acid, geniposide, paeoniflorin, and paeonol in *Dan-zhi-xiao-yao-san* (Figure 1).The neuroprotective effects of Sen A against Cort-induced apoptosis further confirmed by LDH, Hoechst staining (Figure 2B–D). Although SenA has the potential to be a promising neuroprotecitve agent, high dose of the agent (2 mg/ml) was clearly cytotoxic to PC12 cells, reminding the scientist and clinicians that application of SenA in practice should be careful handled.

In addition to Parkinson’s disease, recent studies have shown that the expression of α-syn is associated with depression [24, 27] and overexpression of α-syn may also use a model for depression [24], suggesting the importance of α-syn in depression. In addition, phospho-Ser129 α-syn is the most frequently modifier of α-syn and plays an important role for α-syn induced cell death[28]. Indeed, our results indicate that Cort treated PC12 cells significantly increased α-syn and phospho-Ser129 α-syn levels (Figure 3C). In contrast, SenA attenuated Cort-induced the increase of α-syn and phospho-Ser129 of α-syn levels. PP2A plays a major role in regulating α-syn expression [29]. and α-syn modulates PP2A expression and activities [30]. As expected, Cort-induced the increased of α-syn accompanied by the increase in the expression of PP2A levels (Figure 3C). Importantly, Cort also induced the down-regulation phoshorylation of PP2A (Figure 3C). SenA blocked Cort-induced the changes in the expression of PP2A, p-PP2A, α-syn and p-α-syn levels, suggesting that the neuroprotective effects of SenA may through modulating PP2A and α-syn pathway.

Our results by using PP2A inhibitor OA and activator SPH showed that PP2A is essential for Cort-induced cell apoptosis (Figure 4). The evidence showed that PP2A inhibitor OA coordinate the neuroprotective effects of SenA in Cort-induced cell apoptosis (Figure 4C–D), further confirm that PP2A play a critical role for the neuroprotective effects of SenA. These results is consistent with previous report that PP2Aligand have neuroprotective effects [31].

We next detected the roles of α-syn in the neuroprotective effects of SenA in Cort-induced cell death. Firstly, knockdown and overexpression of α-syn,we found that overexpression of α-syn increased, while knockdown of α-syn decreased cort-induced cell apoptosis (Figure 6E–H), indicated that the expression of α-syn indeed plays an important role for the neuroprotective effects of SenA in Cort-induced cell apoptosis. As phosphorylation of α-syn Ser-129 is essential for α-syn induced cell death[32]. Moreover, PP2A plays an important role in regulating α-syn Ser-129 phosphorylation. To examine how α-syn affects the neuroprotective effects of SenA, we mutated Ser-129 to Ala-129 to prevent its phosphorylation. We demonstrate that α-syn Ala-129 decreased Cort-induced apoptosis (Figure 8B–E) and the neuroprotective effects of SenA can be blocked by α-syn Ser-129 overexpression, suggesting that α-syn Ser-129 is essential for Cort-induced cell apoptosis. Together our data suggest that α-syn levels as well as its Ser-129 phosphorylation is essential for the neuroprotective effects of SenA in Cort-induced cell apoptosis.

In summary, our results, for the first time, demonstrate that SenA plays an important role in protection against Cort-induced cell apoptosis. Importantly, we show that PP2A, α-syn and p-Ser-29-α-syn play critical roles for Cort-induced cell apoptosis. Interestingly, the neuroprotective effects of SenA involved modulating these pathways. Collectively, our results not only demonstrate the importance of PP2A and α-syn in Cort-induced cell death but also revealeda molecular mechanism for the neuroprotective of SenA.

## Conflict of Interests

The authors declare that there is no conflict of interests regarding the publication of this paper.

## Acknowledgment

This work was supported by the Shenzhen Municipal Science and Technology Innovation Council (NO. JCYJ20130328154910812), GZUCM Science Fund for Creative Research Groups (NO. 2016KYTD10), GZUCM Torch Program (XH20140106).

